# An enhanced Eco1 retron editor enables precision genome engineering in human cells without double-strand breaks

**DOI:** 10.1101/2024.08.05.606586

**Authors:** Matthew A. Cattle, Lauren C. Aguado, Samantha Sze, Sanjana Venkittu, Dylan Yueyang Wang, Thales Papagiannakopoulos, Susan Smith, Charles M. Rice, William M. Schneider, John T. Poirier

## Abstract

Retrons are a retroelement class found in diverse prokaryotes that can be adapted to augment CRISPR-Cas9 genome engineering technology to efficiently rewrite short stretches of genetic information in bacteria and yeast. However, efficiency in human cells has been limited by unknown factors. We identified non-coding RNA (ncRNA) instability and impaired Cas9 activity due to 5’ sgRNA extension as key contributors to low retron editor efficiency in human cells. We re-engineered the Eco1 ncRNA to incorporate an exoribonuclease-resistant RNA (xrRNA) pseudoknot from the Zika virus 3’ UTR and devised an RNA processing strategy using Csy4 ribonuclease to minimize 5’ sgRNA extension. This strategy increased steady-state ncRNA levels and rescued sgRNA activity, leading to increased templated repair. This work reveals a previously unappreciated role for ncRNA stability in retron editor efficiency in human cells and presents an enhanced Eco1 retron editor capable of precise genome editing in human cells from a single integrated lentivirus and, in the context of the nCas9 H840A nickase, without creating double-strand breaks.

## INTRODUCTION

Germline and somatic genetic variants play an important role in human health and disease (1-4). CRISPR-Cas9 technology has revolutionized the manipulation of genetic information by enabling RNA-programmable delivery of enzymatic activities to targeted locations in the genome, facilitating new genome editing technologies that can interrogate the relationship between genotype and phenotype in living systems (5). When delivered *via* lentivirus at a low multiplicity of infection (MOI) to ensure a single integration per cell, linked genotype-phenotype relationships can be maintained over multiple cell divisions and the fate of individually edited cells and their progeny can be tracked. This significantly simplifies the scalable parallelization of forward genetic screens, retrieval of rare clones from a population of cells, and somatic editing in genetically engineered models of cancer.

Base editors can install single nucleotide variants (SNVs) with high efficiency but require distinct and exquisitely engineered editing enzymes to generate different types of mutations, limiting their generalizability as genome editing tools (6). In contrast, prime editors offer a greater potential to efficiently introduce a wide spectrum of mutation types, including small insertions; however, each prime editing guide RNA (pegRNA) requires extensive design effort to yield efficient editing, complicating the construction of large pegRNA libraries (7-10). There remains a need for additional methods to complement current genome editing tools that have the properties of broad editing potential, predictable activity, and compatibility with lentiviral delivery.

Retrons are prokaryotic retroelements that play a role in anti-phage defense systems (11-19). Retrons have been adapted as genome editing tools by harnessing their ability to generate homology-directed repair templates *in situ* (20-26). A retron minimally consists of a non-coding RNA (ncRNA) that is reverse transcribed into multicopy single-stranded DNA (msDNA) by its cognate primer-independent reverse transcriptase (27,28). The retron ncRNA contains both the msr stem-loops that are required for recognition and initiation of reverse transcription, as well as the msd segment that contains the reverse transcription template. Arbitrary sequences of interest can be inserted into the msd stem loop, where they are reversed transcribed into RT-DNA (27). Fusion of retron protein and RNA components to their respective CRISPR-Cas9 counterparts results in targeted, templated DNA repair (22). However, the application of retrons as genome editing tools in human cells has been limited due to low efficiency of templated repair and editing outcomes dominated by indels introduced through non-homologous end joining (NHEJ) (23-25).

Previous efforts to improve retron editor efficiency have focused on the cellular abundance of RT-DNA (25,29). Here we determined that retron ncRNA and sgRNA fusion transcripts are unstable in human cells. These unstable ncRNAs exhibit limited templated editing efficiency and Cas9 endonuclease activity, ultimately resulting in diminished activity. Through iterative optimization and rational design, we developed a ncRNA architecture derived from the Eco1 (Ec86) retron with enhanced genome editing efficiency without causing double-strand breaks, reducing unproductive editing outcomes and broadening the utility of retrons as genome editing tools in human cells.

## MATERIAL AND METHODS

### Cell culture

Parental HEK293T cell lines and their derivatives were cultured in high-glucose DMEM with L-glutamine and sodium pyruvate (Gibco) and supplemented with 10% Foundation™ fetal bovine serum (Gemini Bio-Products). K562 cell lines were grown in RPMI 1640 Medium supplemented with L-glutamine (Gibco) and 10% Foundation™ fetal bovine serum (Gemini Bio-Products). Both HEK293T and K562 parental lines were obtained from ATCC. All cell lines were grown at 37ºC and 5% CO_2_. Huh-7.5 cells were maintained in Dulbecco’s modified Eagle’s medium (DMEM; Fisher Scientific, catalog no. 11995065) supplemented with 0.1 mM nonessential amino acids (NEAAs; Fisher Scientific, catalog no. 11140076) and 10% fetal bovine serum (FBS; HyClone Laboratories, lot. #AUJ35777).

### Plasmid cloning

All oligos, and gBlocks used in this study are listed in **Supplemental Table 1**. Plasmids were transformed and propagated in NEB Stable Competent E. coli (New England Biolabs). All ncRNA expressing plasmids were cloned from lentiGuide-Puro (lentiGuide-Puro was a gift from Feng Zhang, Addgene #52963) using the PacI restriction enzyme cloning sites followed by NEBuilder HiFi DNA Assembly (New England Biolabs) of synthesized gBlock™ gene fragments (Integrated DNA Technologies). Cas9-RTwt expression plasmid was cloned from pBZ210 (pBZ210 was a gift from Hunter Fraser, Addgene #170185) by XbaI and PciI double digestion (New England Biolabs) to excise the chimeric sgRNA-retron ncRNA, followed by blunting with DNA Polymerase I Large Klenow fragment (New England Biolabs). The resulting dsDNA blunt ends were then ligated using a T4 ligase (New England Biolabs). Cas9-RTmut was generated by site-directed mutagenesis. TR-Csy4 and TR-Csy4-H29A were gifts from Aravind Asokan (Addgene #80601, Addgene #80602).

### In vitro transcription

T7 expression plasmids were linearized by digestion with EcoRI (New England Biolabs) and then purified by phenol:chloroform extraction followed by ethanol precipitation. A 5μL in vitro transcription reaction was set up with 250ng of linearized plasmid template, 0.375μL of T7 RNA Polymerase (New England Biolabs) and 100mM each of ATP, GTP, CTP, and UTP and then incubated overnight at 37ºC. In vitro transcribed RNA was then purified by phenol:chloroform extraction and ethanol precipitation.

### Plasmid transfection and electroporation

Lipofectamine 3000 was used for all transient transfections of HEK293T-BFP cells. Briefly, 2e5 HEK293T-BFP cells were reverse transfected with 750ng of either Cas9-RTwt or Cas9-RTmut, and 650ng of plasmids expressing the retron editor ncRNA. For all experiments incorporating Csy4 cleavage, 750ng of either Csy4 wild-type or Csy4-H29A was added to the transfection mix, and constructs without a Csy4 recognition site were co-transfected with 750ng pUC19 (New England Biolabs) to balance total transfected mass. Cells were replated in a 12-well plate approximately 16-24 hours after transfection. The Neon NxT Electroporation System was used to electroporate K562-BFP ncRNA-expressing cell lines with Cas9-RT plasmids and one of Csy4, Csy4-H29A, or pUC19 to balance total transfected mass. 5e5 cells were pelleted at 300 RCF for 5 minutes and resuspended in Neon NxT R buffer and then mixed with 2500ng of the Cas9-RT plasmid and 2500ng of TR-Csy4/TR-Csy4-H29A/pUC19 plasmid. Cells were then electroporated using the following parameters in a 10μL Neon NxT tip: 1050V, 20ms pulse width, 2 pulses. Cells were added to 500μL of pre-warmed RPMI media in a 48 well plate after pulsing.

### Flow cytometry

All flow cytometry experiments were done on a BD FACSymphony™ A5 Cell Analyzer. To prepare samples, cells were centrifuged at 350 RCF for 5 minutes and the cell pellets were resuspended in DPBS (Gibco) supplemented with 1% Foundation™ fetal bovine serum (Gemini Bio-Products). Cell populations were analyzed to determine the proportion of BFP and GFP positive cells and each condition was run with three biological replicates. Transfected cells were selected using the mCherry fluorescent reporter on the Cas9-RT expression plasmids. All gates were drawn using untransfected control samples.

### Virus production

T25 flasks were seeded with 2e6 HEK293T cells 24 hours prior to transfection. On the following day, a transfection solution was made up of 200μL OptiMEM (Gibco), 3μg of the lentiviral vector, 2μg of psPAX2, 1μg of pMD2.G, and 12μg of PEI 25K (Polysciences). psPAX2 and pMD2.G were gifts from Didier Trono (Addgene #12260, Addgene #12259). After mixing, the transfection solution was vortexed for 15 seconds, incubated at room temperature for 15 minutes, and added to cells drop wise. The media was replaced 24 hours after transfection. At 72 hours post-transfection the viral supernatant was filtered through a 0.45μm syringe filter and immediately stored at -80ºC.

### Lentivirus transduction

The HEK293T-BFP cell line was made by limiting dilution lentivirus transduction of BFP dest clone lentivirus and selected with 6μg/mL blasticidin (Gibco) for one week. BFP dest clone was a gift from Jacob Corn (Addgene #71825). Cells were then single-cell sorted into a 96-well plate for the top 2.5% fluorescent cells. Single-cell clones were expanded and two clonal lines were randomly selected. Low-copy integrated ncRNA HEK293T-BFP cell lines were generated by limiting dilution lentivirus transduction. 5e5 cells were plated in a 6-well plate and limiting volumes of viral supernatant were added to each well, ranging from 50μL to 1μL. 24 hours after the addition of virus, the cells were selected using 2μg/mL puromycin (Gibco) and grown out on puromycin for one week. For every HEK293T-BFP cell line generated, 1μL of virus was sufficient to yield sparse colonies for subsequent outgrowth. Low-copy integrated ncRNA K562-BFP cell lines were generated by limiting dilution spinfection. Briefly, 1e6 K562-BFP cells were plated in a 6-well plate with 2mL of RPMI 1640 media. After plating, 20μL of lentivirus and 2μL of polybrene (Sigma-Aldrich) were added directly to each well and cells were centrifuged at 930 RCF for 2 hours at room temperature. After centrifugation, 2mL of RPMI 1640 media was added to each well and cells were placed back in the incubator. Selection for infected cells started 24 hours after spinfection using 2μg/mL puromycin. The K562-BFP cell line used in this study was a gift from Dr. Chris Richardson.

### Northern blotting

1e6 HEK293T cells were first reverse transfected in a 6-well plate with 1μg of Cas9-RT (either Cas9-RTwt or Cas9-RTmut) and 1μg of ncRNA expressing plasmid using Lipofectamine 3000 according to the manufacturer’s protocol. Cells were incubated for 48h and then total RNA was extracted using TRIzol reagent (Invitrogen) followed by two chloroform washes. The aqueous layer was loaded onto a Monarch Total RNA Miniprep Kit (New England Biolabs) column and purified according to the manufacturer’s protocol. To radiolabel ssDNA probes, 0.15mCi ATP [*γ* -32P] (Perkin Elmer) was mixed with 200ng of the probe and T4 kinase (New England Biolabs) and then incubated at 37ºC for 60 minutes. Labeled probes were then centrifuged for 30 seconds at 16,000 RCF in a MicroSpin G-50 column (Cytiva) and eluted in 80μL of TNES buffer (50mM Tris, 400mM NaCl, 20mM EDTA, 0.5% SDS). Probe sequences are listed in **Supplemental Table 1**. For electrophoretic size separation, 25ng of in vitro transcribed RNA and 7.5μg of total total RNA was loaded per well onto a Novex 10% TBE-Urea 10-well gel (Invitrogen) and run for 5 hours at constant 120V. RNA was then transferred to a Nytran SuPerCharge (Cytiva) nylon membrane using a Bio-Rad Trans-Blot Turbo (Bio-Rad) at 200mA for 60 minutes. Membranes were blocked at 65ºC for 1 hour in 30mL of 6X SSC 7% SDS blocking buffer. After blocking, membranes were incubated overnight at 42ºC with 30μL of the radiolabeled probe. Membranes were washed and exposed for 48h before imagining. For U6 loading controls, membranes were stripped using 0.1% SDS 2mM EDTA stripping buffer followed by three washes and then reprobed using a U6 loading control probe. Loading control membranes were exposed for 24 hours before imaging.

### Quantitative real-time PCR

First, 5e5 HEK293T-BFP ncRNA-expressing cells were reverse transfected with 1μg of Cas9-RTmut plasmid. 24 hours after transfection, cells were treated with 5μg/mL of actinomycin D (Sigma-Aldrich) and then lysed with TRIzol at the specified time points. The TRIzol lysates were washed with chloroform and the aqueous phases were loaded onto RNeasy Mini Kit columns (Qiagen) and washed and eluted according to the manufacturer’s protocol. To remove residual DNA, the purified RNA samples were treated with DNase I (New England Biolabs), followed by heat-inactivation of DNase I at 70ºC. cDNA was made from the purified RNA using the GoScript Reverse Transcription System (Promega) according to the manufacturer’s protocol with 5μL of input RNA for each reaction. qPCR was performed with PowerUp SYBR Green Master Mix and recorded on QuantStudio3. The primers used for qPCR are listed in **Supplementary Table 1**.

### Statistics and reproducibility

All experiments were performed in triplicate. Outcomes here are presented as the mean ± standard deviation. For all editing outcome assays, statistical significance was determined by two-sided, unpaired t tests using the Holm-Šídák method for multiple testing correction. Throughout the manuscript, P-values are reported as follows - *: P<0.05, **: P<0.01, ***: P<0.001, ****: P<0.0001.

## RESULTS

### Eco1 retron editor efficiency is inferior to synthetic ssDNA templates in human cell lines

To investigate the potential causes of low retron editor efficiency in human cells, we employed the well-studied Eco1 retron. A protein-coding plasmid expresses Cas9 fused to a codon optimized Eco1 reverse transcriptase by a 33 amino acid linker and is followed by a T2A ribosome skipping sequence and mCherry (Cas9-RT mCherry). This construct allows tracking of transfected cells by red-orange fluorescence (**Fig. 1A**). To control for background levels of plasmid templated repair, we constructed a catalytically dead Eco1 mutant (DD196NN, RTmut) (30) and confirmed that the mutant is inactive by qPCR of retron RT-DNA after RNA transfection (**Fig. S1)**. Non-coding RNA plasmids were designed to express the Eco1 ncRNA with a 100 nucleotide repair template internal to the msd sequences as previously described (msr-msd) (22,31). Retron ncRNAs or sgRNAs encoded by separate plasmids were expressed from the U6 RNAPIII promoter and terminated by a poly(T) transcription termination signal (sgBFP) (32).

**Figure 1.**
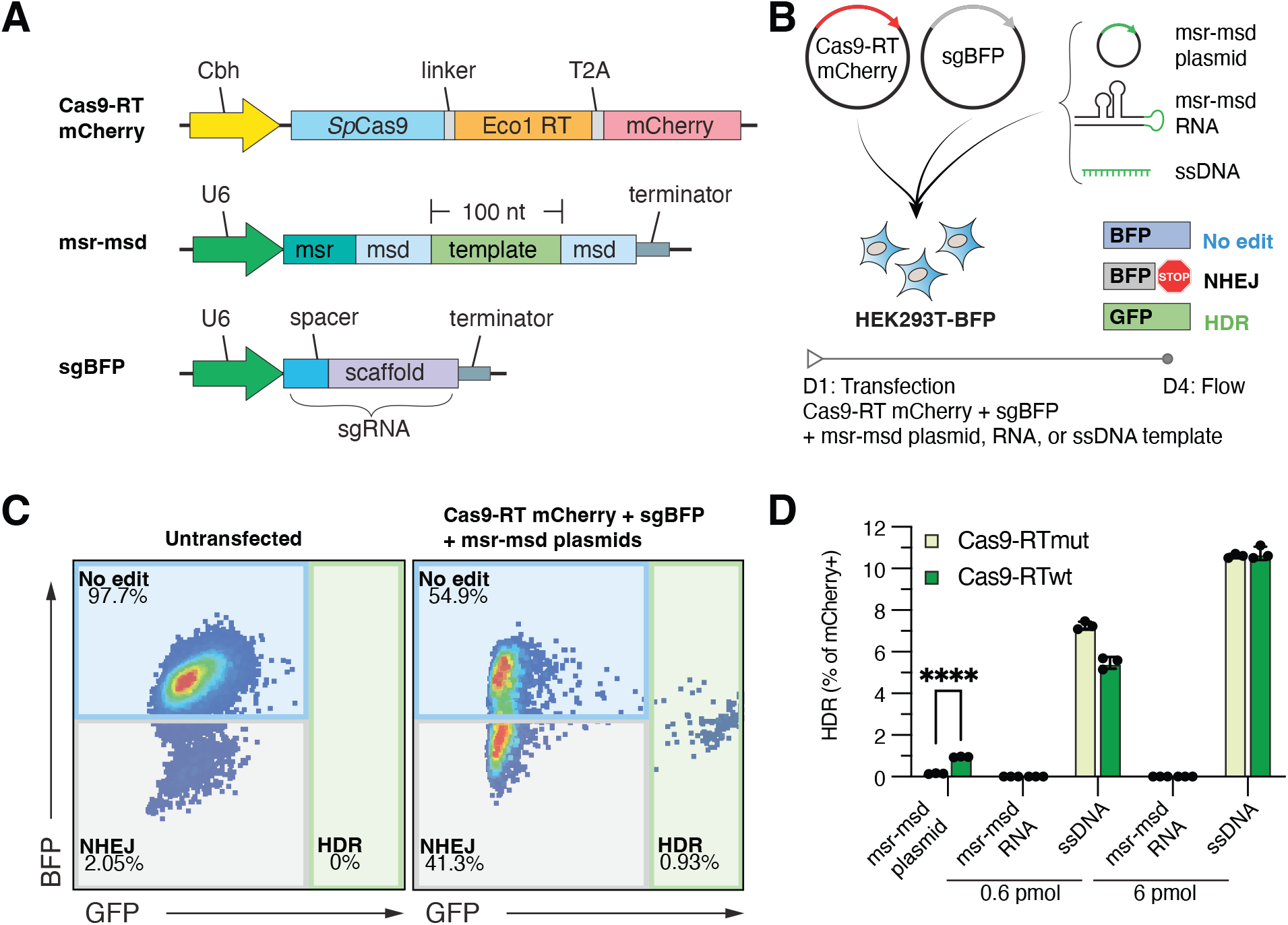
Comparison of Eco1 retron to ssDNA templated repair efficiency in a model genome editing assay. **(A)** Schematic of Cas9-RT mCherry, msr-msd, and sgBFP expression plasmids. **(B)** Experimental setup for transfection-based retron editing. HEK293T-BFP cells were transfected with Cas9-RT mCherry, sgBFP, and one of the indicated repair templates and assessed for editing outcomes by flow cytometry 72h post-transfection. **(C)** Representative flow cytometry plots showing gates used to capture editing outcomes. Gates were drawn using the untransfected control. **(D)** Percentage of GFP positive cells by retron template source at the indicated concentrations. Error bars denote standard error of the mean.

To compare retron editor designs, we measured genome editing rates using an established assay based on the conversion of an integrated cDNA encoding blue fluorescent protein (BFP) to green fluorescent protein (GFP) by missense mutation of the chromophore (31,33). Cas9-RT mCherry and a plasmid encoding an sgRNA targeting the BFP chromophore (sgBFP) were co-transfected with a repair template encoding the mutation conferring GFP-expression and a disruption of the protospacer adjacent motif (PAM) of sgBFP (**Figure 1B**). One of three repair template configurations was used: msr-msd plasmid, in-vitro transcribed (IVT) msr-msd RNA (**Fig. S2)**, or a synthetic 100 nucleotide single-stranded DNA (ssDNA). Editing outcomes were tracked by the relative proportion of BFP+ (no edit), BFP- (NHEJ) or GFP+ (HDR) (**Fig 1C**).

To investigate the contribution of repair template availability, msr-msd RNA and ssDNA were transfected at either 6 pmol, representing a typical quantity of transfected ssDNA repair template (34), or 0.6 pmol, a quantity that we predicted would be limiting. After 72h we quantified editing outcomes (**Fig. 1D**). While the msr-msd plasmid edited the BFP locus to 0.95±0.01%, consistent with prior literature (23-25), transfected msr-msd RNA did not generate any successful editing events. In contrast, the ssDNA conditions resulted in efficient and dose-dependent templated repair of 5.47±0.23% and 10.73±0.15% in the 0.6 pmol and 6 pmol conditions, respectively. These results highlight a positive correlation between ssDNA quantity and editing efficiency and are consistent with the hypothesis that the Eco1 retron produces insufficient suitable homology template in comparison to ssDNA.

### Steady-state levels of retron ncRNAs are limited in human cells

To measure steady-state levels of transfected retron ncRNAs by northern blot, we designed ^32^P radiolabeled ssDNA probes complementary to the sgBFP scaffold sequence present in both the 106 nucleotide sgBFP transcript or the 334 nucleotide full-length fusion transcripts of sgBFP and msr-msd (**Fig. 2A**). Pilot northern blot experiments revealed that co-delivery of Cas9 with sgBFP was required to stabilize sgRNAs (**Fig. S3**). Based on this result, Cas9-RT mCherry plasmid was co-transfected in all subsequent experiments. We transfected plasmids encoding either sgBFP, msr-msd-sgBFP, or sgBFP-msr-msd ncRNAs and extracted total RNA for northern blot analysis (**Fig. 2B**). The abundance of sgBFP, fused in either orientation, to msr-msd ncRNA was lower than sgBFP alone. We also observed that while the U6 snRNA loading control showed no evidence of RNA degradation, there was a consistent pattern of intermediate length ncRNA products that were not observed in *in vitro* transcribed RNA samples. We hypothesized that these intermediate-length products originated through degradation by endogenous nucleases rather than early transcription termination, as they were present even when probing the 3’ end of the ncRNA.

**Figure 2.**
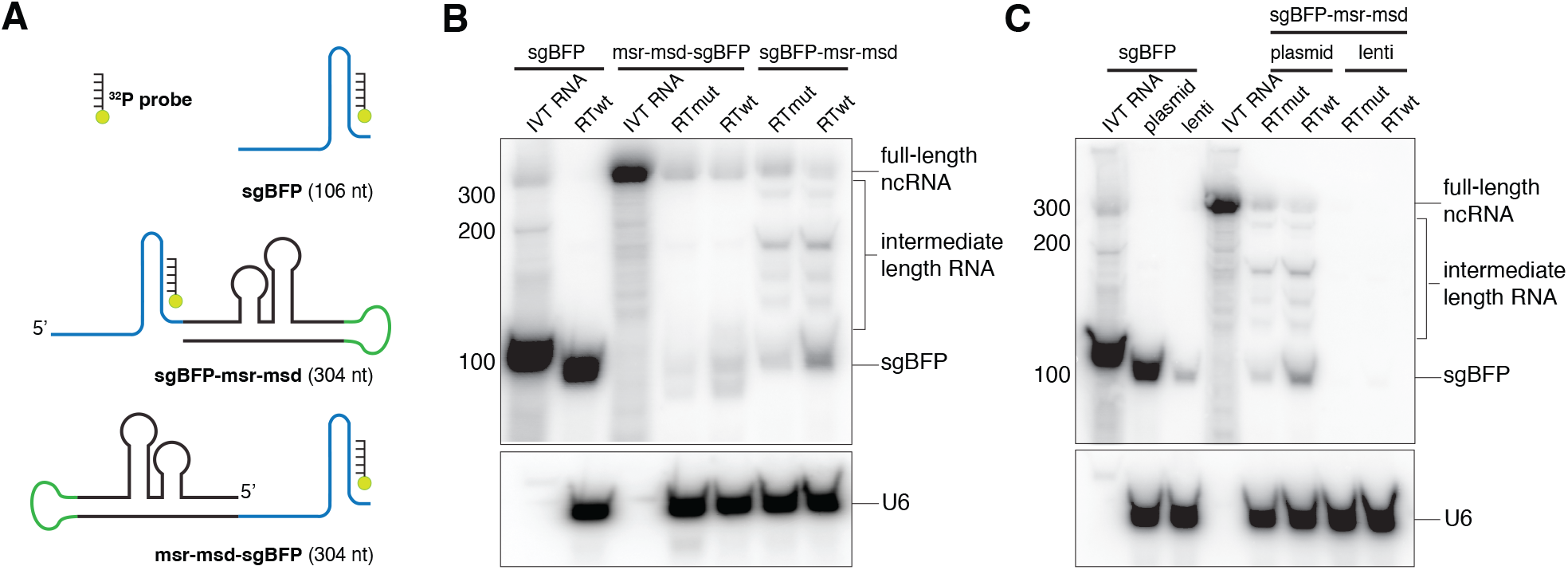
Analysis of ncRNA integrity and quantity when expressed from plasmid and lentiviral vectors. **(A)** Schematic of both 3’ and 5’ sgBFP orientations depicting location of the radiolabeled probe in the sgRNA scaffold sequence. **(B)** Northern blot of retron RNAs expressed from transfected plasmids for sgBFP, msr-msd-sgBFP, and sgBFP-msr-msd. **(C)** Northern blot showing RNA levels of sgBFP or sgBFP-msr-msd when expressed from either transient transfection of a plasmid or low copy integration of a lentivirus vector.

ncRNA expression in a low-copy lentivirus integration cell line with a sgBFP-msr-msd ncRNA construct was compared to RNA expression from a cell line with low-copy integrations of sgBFP alone (**Fig. 2C**). We detected a band corresponding to full-length sgBFP when expressed from a transfected plasmid and from an integrated locus; however, we were unable to detect the full-length sgBFP-msr-msd ncRNA when integrated despite being expressed from the same promoter and plasmid backbone as sgBFP. Taken together, these results were indicative of ncRNA degradation by endogenous nucleases.

### Efficient retron editing favors ncRNA architectures encoding sgRNAs with native 5’ ends

To determine which elements of ncRNA architecture most strongly affected editing efficiency, we systematically tested different ncRNA designs by transient transfection of retron editor plasmids. Initially we compared editing efficiencies of *cis* or *trans* expression of the sgRNA and retron template. In this case, *cis* refers to a fusion transcript of sgRNA and retron msr-msd components while *trans* refers to expression of sgRNA and retron msr-msd as two separate transcripts. For editing in the context of a fusion RNA, we were interested in how orientation of the sgRNA relative to msr-msd might impact retron editing efficiency (i.e., msr-msd-sgBFP vs. sgBFP-msr-msd).

We observed that with Cas9-RTwt the msr-msd-sgBFP orientation showed no significant increase in GFP+ cells as compared to Cas9-RTmut (0.36±0.07% versus 0.37±0.04%), indicating that the GFP+ cells likely result from plasmid-templated repair in this context (**Fig. 3A**). However, *trans* expression of the sgRNA and msr-msd from two different plasmids (sgBFP + msr-msd) and an sgBFP-msr-msd orientation had a marginal but statistically significant increase in editing efficiency (0.43±0.08% and 0.64±0.06% respectively). These results demonstrate that the overall architecture of the retron ncRNA favors an sgRNA with a native 5’ end.

**Figure 3.**
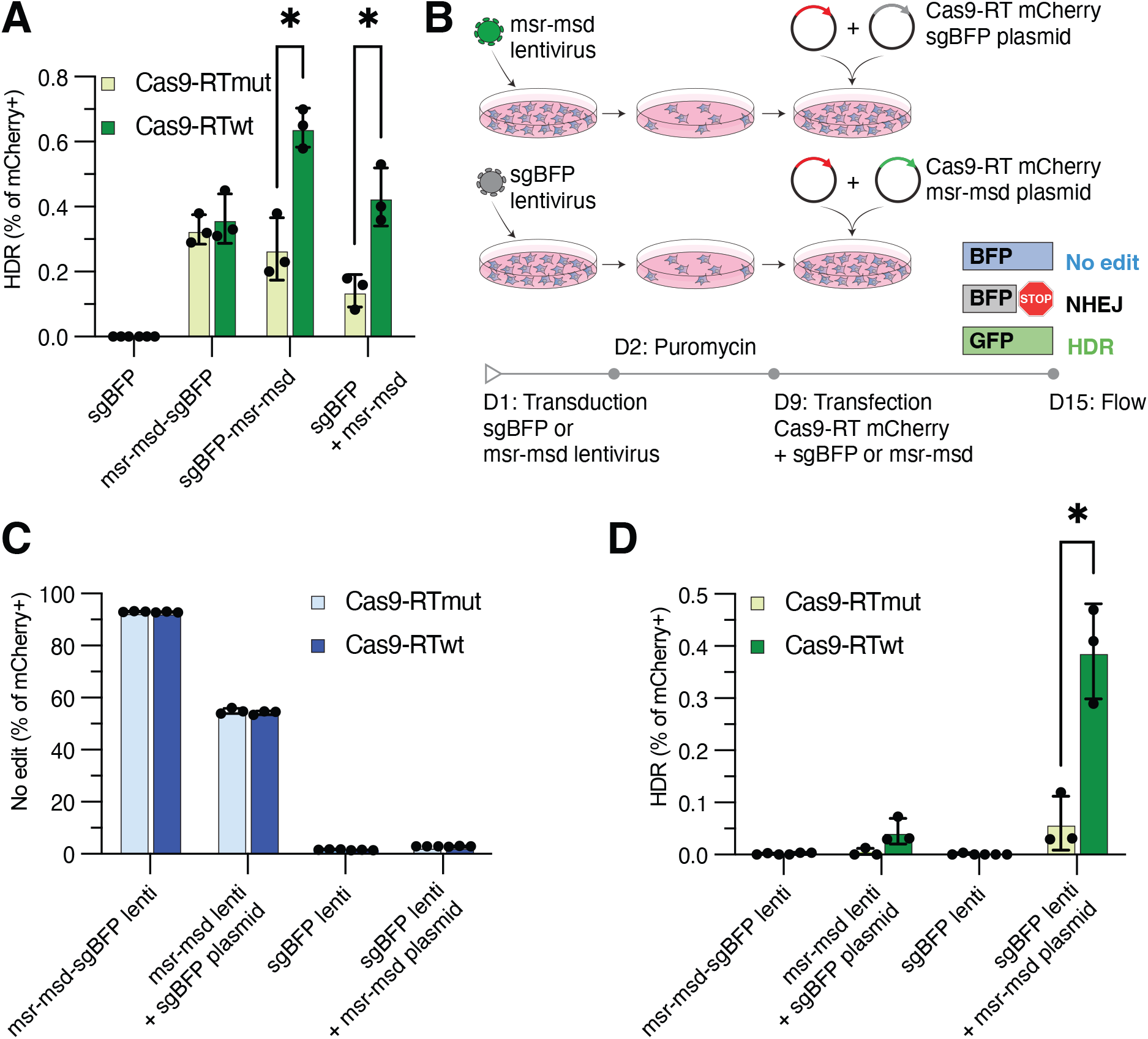
Genome editing efficiency of retron editor architectures and delivery strategies. **(A)** HDR editing outcomes from transfection-based editing assay as indicated by percentage of GFP+ cells. **(B)** Experimental setup for integrated ncRNA retron editing. **(C)** NHEJ or **D)** HDR editing outcomes in integrated or transfected contexts. Error bars denote standard error of the mean.

The importance of the relative abundance of the sgRNA and retron RNA template is an aspect of retron-mediated genome editing that has not previously been explored. We exploited our finding that retron ncRNA components are functional in *trans* to interrogate the relationship between ncRNA and sgRNA abundance and retron editing efficiency under conditions where one RNA is limiting. We packaged either sgBFP or msr-msd as lentivirus, and transduced HEK293T-BFP cells at low MOI to bias toward low copy number. Subsequently, we transfected the cognate component to induce overexpression relative to the low copy integrated component (**Fig. 3B**). When integrated and expressed with Cas9-RT mCherry, sgBFP disrupted BFP at a mean frequency of 98.28±0.14%, while conversely, the msr-msd-sgBFP fusion was unable to disrupt BFP at a frequency greater than 6.93±0.15% (**Fig. 3C**). Efficient BFP disruption was restored when sgBFP was expressed with msr-msd template in *trans* either by transfection or integration. Overall, templated editing efficiency from integrated constructs was less than 0.1% except when sgBFP was integrated in the genome and the msr-msd template was overexpressed by plasmid transfection (**Fig. 3D**). Under these conditions, templated repair overwhelmingly originates from RT-DNA, evidenced by a significant increase in GFP+ cells with Cas9-RTwt as compared to Cas9-RTmut which establishes the baseline level of plasmid templated repair. These results suggest that the msr-msd template levels could be limiting in human cells and that retron-mediated editing efficiency might be improved by increasing the abundance of the ncRNA. These results also indicate that fusion of the sgRNA to the retron msr-msd inhibits Cas9 nuclease activity, potentially by interfering with efficient RNP assembly due to misfolding or steric hindrance (35).

### xrRNA pseudoknot rescues Eco1 retron editor ncRNA steady-state levels

Free 5’ and 3’ RNA ends are substrates for exoribonucleases and RNA sensing pathways (36). We hypothesized that protecting free msr-msd ends would stabilize and increase msr-msd abundance and improve editing efficiency. Since the sgRNA is already protected by co-expression with Cas9, we focused on designs aimed at stabilizing the 5’ end of the msr-msd-sgRNA fusion orientation. This orientation also places the repair template proximal to the sgRNA spacer sequence, which is advantageous for molecular cloning. We tested three strategies of RNA protection found in nature: RNA circularization (37), 3’ polyadenylation (38), and RNA pseudoknots (39). To generate a circular RNA (circRNA) construct, we took advantage of flanking tRNA ligation sequences as endogenous tRNA ligase activity has previously been used to extend sgRNA half-life in cells (40).

Polyadenylation of RNAPIII-expressed SINE transcripts has been achieved by the addition of a canonical AATAAA polyA signal to the 3’ end of the transcript, and we reasoned that we could similarly take advantage of the stability provided by a polyA tail (38). Finally, we tested three additional approaches incorporating stability-enhancing RNA pseudoknots on the 5’ end that all have been previously used to stabilize pegRNA transcripts and improve prime editing efficiency (9,41).

We tested five distinct ncRNA plasmids with the intention of circularizing, polyadenylating, or pseudoknot-protecting the retron editor ncRNA (**Fig. 4A**) and tested the degree to which each construct stabilized Eco1 retron editor transcripts within the cellular environment. Total RNA was extracted from HEK293T-BFP cells co-transfected with the different msr-msd plasmids and either Cas9-RT or Cas9-RTmut. Steady state RNA levels were probed by northern blot as described above (**Fig. 4B**).

**Figure 4.**
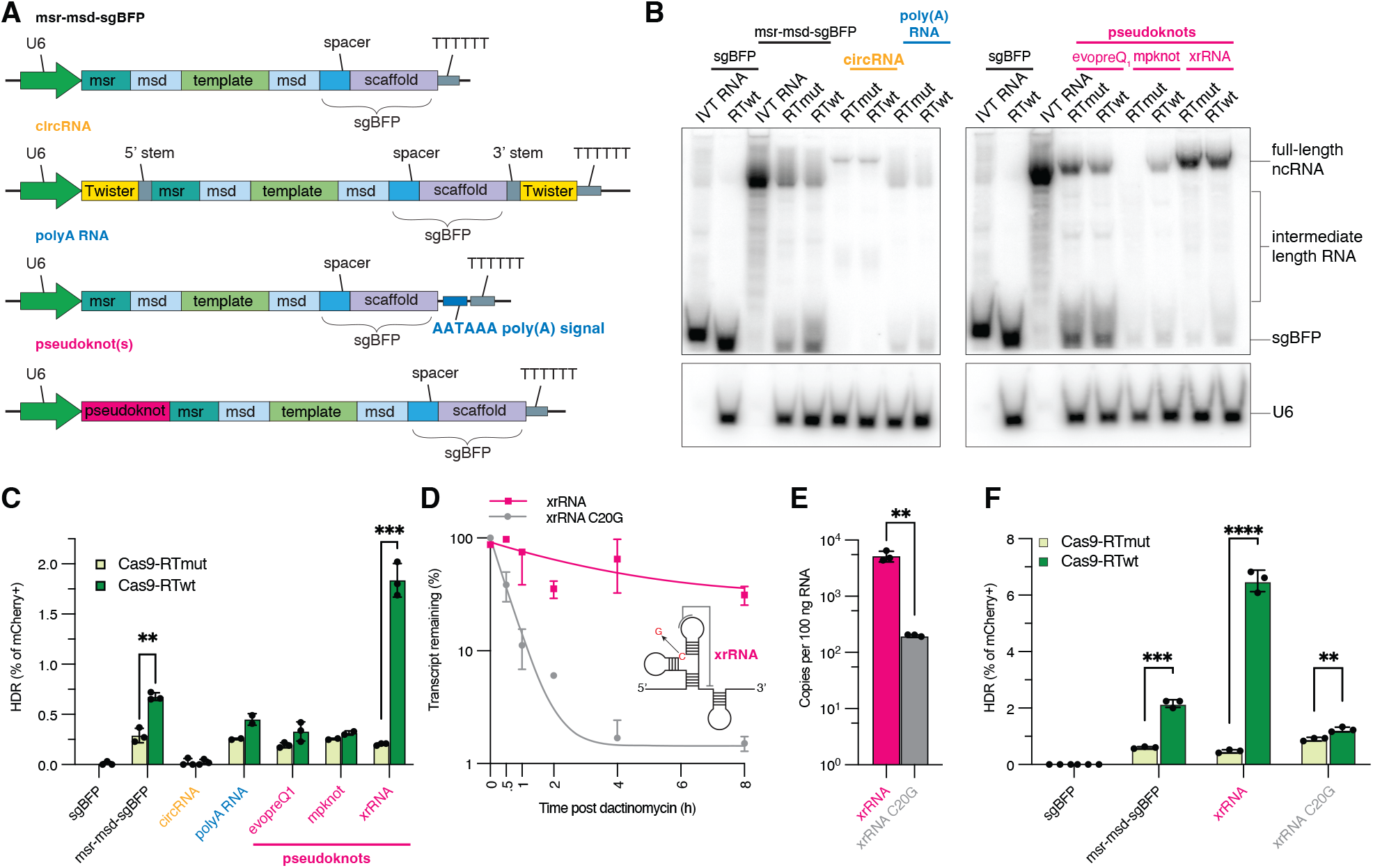
Evaluation of strategies to rescue retron ncRNA expression. **(A)** Diagrams of retron editor ncRNA designs incorporating stabilizing elements. Top: circRNA design using Twister ribozyme sequences and tRNA stem forming ligation sequences. Middle: polyA RNA design incorporating a polyA AATAAA sequence before the RNAP III termination sequence. Bottom: ncRNA designs with a protective pseudoknot on the 5’ end before the retron template. **(B)** Northern blots of retron editor ncRNAs incorporating stabilizing elements. Radiolabeled probe targeted the scaffold sequence of the sgRNA. **(C)** GFP editing outcomes for ncRNA constructs incorporating stability-enhancing elements. **(D)** RNA abundance at the indicated time points after dactinomycin treatment (inset, xrRNA C20G mutation in context). **(E)** Steady-state RNA abundance in copies per 100 ng input RNA. **(F)** GFP editing outcomes with functional xrRNA and misfolded xrRNA C20G. All error bars denote standard error of the mean.

Next, we compared editing efficiencies of these different stability enhancing ncRNAs to msr-msd-sgBFP. When transfected with Cas9-RTwt, the xrRNA-msr-msd-sgBFP reached GFP editing rates as high as 1.83±0.16%, representing a nearly 3-fold improvement over the native unprotected ncRNA (**Fig. 4C**). No other ncRNAs tested improved templated editing rates, while the circular RNA design exhibited reduced performance due to significantly impaired Cas9 activity, resulting in <0.1% editing efficiency.

Since the xrRNA pseudoknot evolved in nature to block nuclease-mediated degradation of an RNA genome (42,43) we hypothesized that a similar mechanism could explain the increase in RNA abundance in this context. To test whether exonuclease resistance is required for the increase in editing efficiency imparted by the xrRNA, we introduced a single point mutation (C20G) known to disrupt the pseudoknot three-way junction and decrease exonuclease resistance while maintaining overall secondary structure of the xrRNA stem loops (44,45) (**Fig. 4D**, inset). To assess differences in half-life, we treated HEK293T cells expressing xrRNA or xrRNA C20G transcripts with dactinomycin to inhibit transcription and collected samples at multiple timepoints over 8 hours for analysis. Total RNA was extracted and the abundance of each transcript was quantitated by reverse transcription followed by real-time quantitative PCR. The xrRNA C20G transcript rapidly decreased in abundance by single phase exponential decay (R^2^=0.99) with an apparent half-life of 20 minutes (16.8–24 95% CI) (**Fig. 4D**). In contrast, the xrRNA transcript decreased monotonically in a sub-exponential manner, reaching approximately one third of the initial quantity by 8 hours. The increased stability of the xrRNA transcript compared to the xrRNA C20G transcript coincided with a steady-state abundance ∼27 fold greater (**Fig. 4E**). Functionally, the C20G point mutation abolished any increase in editing efficiency, supporting exonuclease resistance as the aspect of the xrRNA-msd-msd-sgBFP ncRNA that enhances retron editing efficiency (**Fig. 4F)**.

### Csy4 cleavage and pseudoknot protection enable Eco1 retron editing without double-strand breaks

Despite the improvements in xrRNA-msd-msr-sgBFP abundance and editing efficiency in the context of transient plasmid transfection, this design showed low BFP disruption rates of 13.06±0.28% and an absence of templated repair as a likely consequence of this modification (**Fig. S4A**). Based on prior reports that sgRNA activity is reduced by 5’ extensions (35) and given our results demonstrating reduced BFP disruption in the context of 5’ sgBFP fusions, we reasoned that Cas9 activity might be if the msr-msd RNA was enzymatically cleaved from the sgRNA after transcription, thereby liberating the sgRNA and minimizing 5’ sgRNA extension.

We explored multiple strategies to cleave the msr-msd-sgBFP including tRNA sequences that have natural processing of their 5’ and 3’ ends by cellular RNases (46), as well as the Csy4 endoribonuclease that cleaves at a specific 20 nt stem loop sequence (47). We tested three processable retron editor ncRNAs: two tRNA sequences and one Csy4-cleavable transcript. tRNAPro represents a full-length prolyl tRNA sequence between the template and sgRNA, while dC55G incorporates an engineered prolyl tRNA with reduced internal promoter activity and improved RNA processing (48). When co-transfected with Cas9-RT and a plasmid expressing wild-type Csy4, the Csy4-cleavable retron editor ncRNA (msr-msd-csy4-sgBFP) showed marginal improvements in RT-dependent templated repair as compared to an unprocessed ncRNA.(**Fig. S4B**) This improvement is Csy4 cleavage dependent, as increased editing was not observed when co-transfected with a plasmid expressing a nuclease deficient Csy4 mutant (Csy4-H29A). Processing the sgRNA to minimize 5’ extension serves to further improve editing efficiency of the Eco1 retron editor ncRNA.

To overcome the limitations of low steady-state RNA and limited Cas9 nuclease activity due to fusion ncRNA retron editor architectures, we combined the strategies of pseudoknot protection and Csy4 cleavage into a single ncRNA construct: xrRNA-msr-msd-csy4-sgBFP (**Fig. 5A**),wherein an xrRNA pseudoknot is grafted onto the 5’ end of the msr-msd template and a Csy4 recognition site is interposed between the template and the sgRNA. We also tested xrRNA-msr-msd-evopreQ-Csy4-sgBFP, which incorporates an additional pseudoknot on the 3’ end of the template that is exposed after Csy4 cleavage for additional nuclease protection of the mature msr-msd.

**Figure 5.**
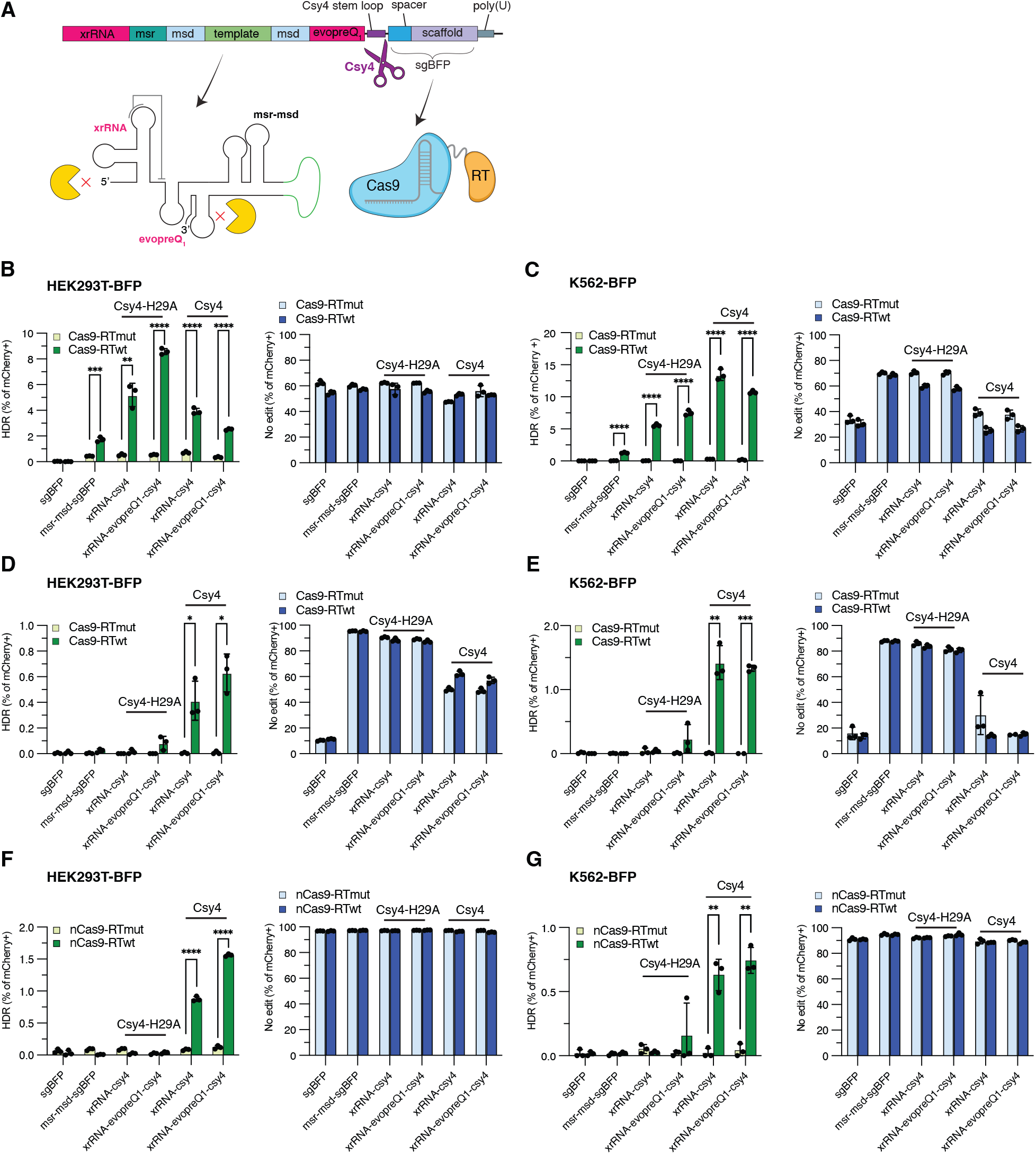
Post-transcriptional processing of ncRNAs to restore a native sgRNA 5’ end. **(A)** Diagram of pseudoknot protected and Csy4-protected retron editor ncRNAs. xrRNA-csy4 contains a single xrRNA pseudoknot on the 5 ‘end of the retron template, while xrRNA-evopreQ_1_-csy4 contains two pseudoknots that flank the retron template for both 5’ and 3’ protection after Csy4 processing. **(B)** Editing efficiency of transfected processed retron editor ncRNAs in HEK293T-BFP cells and in **(C)** K562-BFP cells. **(D)** Editing efficiency in low-copy integrated HEK293T-BFP cells and in **(E)** K562-BFP cells. **(F)** Editing efficiency using nCas9-H840A in HEK293T-BFP cells and in **(G)** K562-BFP cells.

In HEK293T-BFP cells transiently transfected with the enhanced Eco1 retron editor, we observed a 2.26-fold increase in templated editing (**Fig. 5B**) while in the K562-BFP chronic myelogenous leukemia cell line transfected with the improved Eco1 retron editor, we observed a 10.63-fold increase in templated editing (**Fig. 5C**). Contrary to our initial expectations and our findings in K562-BFP cells, we observed consistently greater templated editing in HEK293T-BFP cells co-transfected with Csy4-H29A versus wild-type Csy4. This result led us to consider whether reduced sgRNA efficiency could be beneficial in certain contexts by limiting the occurrence of dead-end NHEJ outcomes prior to RT-DNA formation. When tested in low-copy HEK293T-BFP cell lines, the enhanced Eco1 xrRNA-msr-msd-evopreQ-csy4-sgBFP construct led to a 27-fold increase in templated editing when compared to the native Eco1 retron editor ncRNA (**Fig. 5D**). In a similar experiment conducted in K562-BFP cells, we observed BFP disruption frequencies of templated editing rates of 1.33±0.03% versus <0.01% using the unoptimized ncRNA construct (**Fig. 5E**).

Finally, we hypothesized that this improved editing efficiency could enable editing without introducing double strand breaks. We mutated the Cas9-RT mCherry vectors to encode the H840A nickase mutant (nCas9) and discovered that in HEK293T-BFP cells (**Fig. 5F**) and in K562-BFP cells (**Fig. 5G**), it was possible to achieve precise genome editing without evident BFP disruption. Amplicon sequencing confirmed indel frequency of ∼5% (**Fig. S5)**. Editing in the context of nCas9-RT was entirely dependent on a functional RT, 5’ xrRNA, and Csy4 processing. In the context of HEK293T-BFP cells was significantly improved by 3’ evopreQ_1_ end protection.

## DISCUSSION

Retron-mediated genome editing is an emerging biotechnology that complements existing tools for precision rewriting of genomic DNA. Retrons generate short ssDNA *in situ* for templated repair, leading to predictable editing outcomes and enabling applications that require large and complex DNA edits in diverse sequence contexts. Previously, retron editors have been used to introduce missense mutations in human cells but their utility is limited by poor efficiency when compared to orthologous systems expressed in bacteria and yeast (20,22).

In this study, we identified unstable Eco1 ncRNA as a factor influencing low efficiency of templated genome editing in human cells. To overcome this limitation, we engineered a series of retron ncRNA variants to incorporate stability-enhancing secondary structures including a nuclease-resistant RNA pseudoknot from the Zika virus 3’ UTR that robustly rescued steady-state RNA levels in the context of transfected and stably integrated constructs. We further identified impaired Cas9 endonuclease activity in the context of a chimeric msr-msd-sgRNA fusion transcript and evaluated strategies to restore the native ends of both RNAs. Enhanced editing required the Csy4 endoribonuclease to enzymatically process the sgRNA from the retron ncRNA components, which rescued Cas9 activity when the RNA components were expressed as a single transcript. Combining these approaches enabled efficient precision genome editing by transient transfection, editing from a lentiviral vector integrated at low copy, and editing without introducing double strand breaks when co-transfected with the Cas9 H840A nickase (nCas9-RT) mutant (49).

The utility of genome editing technologies based on CRISPR-Cas systems has been expanded by fusing various enzymes to Cas9 to recruit enzymatic activities to target DNA. In the case of prime editing, this also requires extending the sgRNA from the 3’ end to encode information for reverse transcriptase priming and the recombination template, which is referred to as a pegRNA (7). The 3’ components of a pegRNA are exposed to degradation by endogenous nucleases (41). Previous studies have identified approaches to protect the 3’ ends of pegRNAs including extension by RNA secondary structures and overexpression of the small RNA-binding exonuclease protection factor La (50). In the present study, stabilization of the sgRNA was not required as it is protected when co-expressed with Cas9-RT; instead, we employed a strategy to protect both the 5’ and 3’ ends of the Eco1 ncRNA, which, in this context, significantly outperformed the native ncRNA. Recent studies have shown that 3’ truncated pegRNAs are preferentially loaded into Cas9 and that pegRNAs with strong 5’ and 3’ complementarity could circularize into a structure similar to retron ncRNA:sgRNA fusions (51,52). These findings are consistent with our data that ncRNA:sgRNA fusions and circRNAs are relatively inefficient as Cas9 guide RNAs.

Further improvements in efficiency may also be realized through rational design, directed evolution, or metagenomic studies aimed at increasing the efficiency and processivity of reverse transcription in human cells or by increasing the suitability of the msDNA structure as a substrate for templated repair. Since the msDNA arises from the ncRNA, we expect RT improvements to complement ncRNA stabilization. We believe that it will be possible to enhance the genome editing utility of retron editors by: using PAM-expanded Cas variants, harnessing the intrinsic RNase activity of Cas12a to streamline RNA processing (53), and/or implementing alternative promoter strategies to overcome the processivity limitations of the U6 promoter (32).

This study investigated the limitations of Eco1 retron-mediated genome editing in human cells. Through systematic modifications and comparative assessments, we have identified key factors influencing the efficiency of a model retron editor. Incorporating RNA-stabilizing pseudoknots and Csy4 cleavage yielded an enhanced Eco1 retron editor capable of writing sequences to the genome without double-strand breaks. Genome editing tools that rely on double-strand breaks to introduce templated missense mutations are limited by a general preference for NHEJ as a repair mechanism, which biases editing outcomes away from the intended edit (54,55). This advance in retron editing resolves one of the most significant limitations of the technology by reducing the frequency of unproductive editing outcomes.

## Supporting information

Combined Supplemental Materials

## SUPPLEMENTARY DATA

Supplementary Data are available at NAR online.

## ACKNOWLEDGEMENTS

We thank our colleagues Drs. Jef Boeke, Benjamin tenOever, and Timothee Lionnet as well as the members of the Rice and Poirier labs for their helpful discussions. Some instrumentation was provided by NYU Langone’s Small Instrument Fleet.

## REFERENCES

Claussnitzer, M., Cho, J.H., Collins, R., Cox, N.J., Dermitzakis, E.T., Hurles, M.E., Kathiresan, S., Kenny, E.E., Lindgren, C.M., MacArthur, D.G. et al./person-group>. (2020) A brief history of human disease genetics. Nature, 577, 179–189.

Sud, A., Kinnersley, B. and Houlston, R.S. (2017) Genome-wide association studies of cancer: current insights and future perspectives. Nat Rev Cancer, 17, 692–704.

Consortium, I.T.P.-C.A.o.W.G. (2020) Pan-cancer analysis of whole genomes. Nature, 578, 82–93.

Dagogo-Jack, I. and Shaw, A.T. (2018) Tumour heterogeneity and resistance to cancer therapies. Nat Rev Clin Oncol, 15, 81–94.

Anzalone, A.V., Koblan, L.W. and Liu, D.R. (2020) Genome editing with CRISPR-Cas nucleases, base editors, transposases and prime editors. Nat Biotechnol, 38, 824–844.

Rees, H.A. and Liu, D.R. (2018) Base editing: precision chemistry on the genome and transcriptome of living cells. Nat Rev Genet, 19, 770–788.

Anzalone, A.V., Randolph, P.B., Davis, J.R., Sousa, A.A., Koblan, L.W., Levy, J.M., Chen, P.J., Wilson, C., Newby, G.A., Raguram, A. et al./person-group>. (2019) Search-and-replace genome editing without double-strand breaks or donor DNA. Nature, 576, 149–157.

Chen, P.J., Hussmann, J.A., Yan, J., Knipping, F., Ravisankar, P., Chen, P.F., Chen, C., Nelson, J.W., Newby, G.A., Sahin, M. et al./person-group>. (2021) Enhanced prime editing systems by manipulating cellular determinants of editing outcomes. Cell, 184, 5635–5652 e5629.

Nelson, J.W., Randolph, P.B., Shen, S.P., Everette, K.A., Chen, P.J., Anzalone, A.V., An, M., Newby, G.A., Chen, J.C., Hsu, A. et al./person-group>. (2022) Engineered pegRNAs improve prime editing efficiency. Nat Biotechnol, 40, 402–410.

Gould, S.I., Wuest, A.N., Dong, K., Johnson, G.A., Hsu, A., Narendra, V.K., Atwa, O., Levine, S.S., Liu, D.R. and Sanchez Rivera, F.J. (2024) High-throughput evaluation of genetic variants with prime editing sensor libraries. Nat Biotechnol.

Millman, A., Bernheim, A., Stokar-Avihail, A., Fedorenko, T., Voichek, M., Leavitt, A., Oppenheimer-Shaanan, Y. and Sorek, R. (2020) Bacterial Retrons Function In Anti-Phage Defense. Cell, 183, 1551–1561 e1512.

Gonzalez-Delgado, A., Mestre, M.R., Martinez-Abarca, F. and Toro, N. (2021) Prokaryotic reverse transcriptases: from retroelements to specialized defense systems. FEMS Microbiol Rev, 45.

Palka, C., Fishman, C.B., Bhattarai-Kline, S., Myers, S.A. and Shipman, S.L. (2022) Retron reverse transcriptase termination and phage defense are dependent on host RNase H1. Nucleic Acids Res, 50, 3490–3504.

Bobonis, J., Mitosch, K., Mateus, A., Karcher, N., Kritikos, G., Selkrig, J., Zietek, M., Monzon, V., Pfalz, B., Garcia-Santamarina, S. et al./person-group>. (2022) Bacterial retrons encode phage-defending tripartite toxin-antitoxin systems. Nature, 609, 144–150.

Sasaki, T., Takita, S., Fujishiro, T., Shintani, Y., Nojiri, S., Yasui, R., Yonesaki, T. and Otsuka, Y. (2023) Phage single-stranded DNA-binding protein or host DNA damage triggers the activation of the AbpAB phage defense system. mSphere, 8, e0037223.

Carabias, A., Camara-Wilpert, S., Mestre, M.R., Lopez-Mendez, B., Hendriks, I.A., Zhao, R., Pape, T., Fuglsang, A., Luk, S.H., Nielsen, M.L. et al./person-group>. (2024) Retron-Eco1 assembles NAD(+)-hydrolyzing filaments that provide immunity against bacteriophages. Mol Cell, 84, 2185–2202 e2112.

Wang, Y., Wang, C., Guan, Z., Cao, J., Xu, J., Wang, S., Cui, Y., Wang, Q., Chen, Y., Yin, Y. et al./person-group>. (2024) DNA methylation activates retron Ec86 filaments for antiphage defense. Cell Rep, 43, 114857.

George, J.T., Burman, N., Wilkinson, R.A., de Silva, S., McKelvey-Pham, Q., Buyukyoruk, M., Dale, A., Landman, H., Graham, A., DeLuca, S.Z. et al./person-group>. (2025) Structural basis of antiphage defense by an ATPase-associated reverse transcriptase. bioRxiv.

Mayo-Munoz, D., Li, H., Mestre, M.R. and Pinilla-Redondo, R. (2025) The role of noncoding RNAs in bacterial immunity. Trends Microbiol, 33, 208–222.

Farzadfard, F. and Lu, T.K. (2014) Synthetic biology. Genomically encoded analog memory with precise in vivo DNA writing in living cell populations. Science, 346, 1256272.

Simon, A.J., Ellington, A.D. and Finkelstein, I.J. (2019) Retrons and their applications in genome engineering. Nucleic Acids Res, 47, 11007–11019.

Sharon, E., Chen, S.A., Khosla, N.M., Smith, J.D., Pritchard, J.K. and Fraser, H.B. (2018) Functional Genetic Variants Revealed by Massively Parallel Precise Genome Editing. Cell, 175, 544–557 e516.

Kong, X., Wang, Z., Zhang, R., Wang, X., Zhou, Y., Shi, L. and Yang, H. (2021) Precise genome editing without exogenous donor DNA via retron editing system in human cells. Protein Cell, 12, 899–902.

Zhao, B., Chen, S.A., Lee, J. and Fraser, H.B. (2022) Bacterial Retrons Enable Precise Gene Editing in Human Cells. CRISPR J, 5, 31–39.

Lopez, S.C., Crawford, K.D., Lear, S.K., Bhattarai-Kline, S. and Shipman, S.L. (2022) Precise genome editing across kingdoms of life using retron-derived DNA. Nat Chem Biol, 18, 199–206.

Tang, S. and Sternberg, S.H. (2023) Genome editing with retroelements. Science, 382, 370–371.

Inouye, M. and Inouye, S. (1992) Retrons and multicopy single-stranded DNA. J Bacteriol, 174, 2419–2424.

Mestre, M.R., Gonzalez-Delgado, A., Gutierrez-Rus, L.I., Martinez-Abarca, F. and Toro, N. (2020) Systematic prediction of genes functionally associated with bacterial retrons and classification of the encoded tripartite systems. Nucleic Acids Res, 48, 12632–12647.

Bhattarai-Kline, S., Lear, S.K., Fishman, C.B., Lopez, S.C., Lockshin, E.R., Schubert, M.G., Nivala, J., Church, G.M. and Shipman, S.L. (2022) Recording gene expression order in DNA by CRISPR addition of retron barcodes. Nature, 608, 217–225.

Sharma, P.L., Nurpeisov, V. and Schinazi, R.F. (2005) Retrovirus reverse transcriptases containing a modified YXDD motif. Antivir Chem Chemother, 16, 169–182.

Richardson, C.D., Ray, G.J., DeWitt, M.A., Curie, G.L. and Corn, J.E. (2016) Enhancing homology-directed genome editing by catalytically active and inactive CRISPR-Cas9 using asymmetric donor DNA. Nat Biotechnol, 34, 339–344.

Gao, Z., Herrera-Carrillo, E. and Berkhout, B. (2018) Delineation of the Exact Transcription Termination Signal for Type 3 Polymerase III. Mol Ther Nucleic Acids, 10, 36–44.

DeWitt, M.A., Corn, J.E. and Carroll, D. (2017) Genome editing via delivery of Cas9 ribonucleoprotein. Methods, 121-122, 9–15.

Chen, F., Pruett-Miller, S.M. and Davis, G.D. (2015) Gene editing using ssODNs with engineered endonucleases. Methods Mol Biol, 1239, 251–265.

Kawamata, M., Suzuki, H.I., Kimura, R. and Suzuki, A. (2023) Optimization of Cas9 activity through the addition of cytosine extensions to single-guide RNAs. Nat Biomed Eng, 7, 672–691.

Wilusz, C.J., Wormington, M. and Peltz, S.W. (2001) The cap-to-tail guide to mRNA turnover. Nat Rev Mol Cell Biol, 2, 237–246.

Unti, M.J. and Jaffrey, S.R. (2024) Highly efficient cellular expression of circular mRNA enables prolonged protein expression. Cell Chem Biol, 31, 163–176 e165.

Ustyantsev, I.G., Tatosyan, K.A., Stasenko, D.V., Kochanova, N.Y., Borodulina, O.R. and Kramerov, D.A. (2020) [Polyadenylation of Sine Transcripts Generated by RNA Polymerase III Dramatically Prolongs Their Lifetime in Cells]. Mol Biol (Mosk), 54, 78–86.

Staple, D.W. and Butcher, S.E. (2005) Pseudoknots: RNA structures with diverse functions. PLoS Biol, 3, e213.

Wang, M., Xu, J., Meng, J. and Huang, X. (2022) Synthetic Circular gRNA Mediated Biological Function of CRISPR-(d)Cas9 System. Front Cell Dev Biol, 10, 863431.

Zhang, G., Liu, Y., Huang, S., Qu, S., Cheng, D., Yao, Y., Ji, Q., Wang, X., Huang, X. and Liu, J. (2022) Enhancement of prime editing via xrRNA motif-joined pegRNA. Nat Commun, 13, 1856.

Pijlman, G.P., Funk, A., Kondratieva, N., Leung, J., Torres, S., van der Aa, L., Liu, W.J., Palmenberg, A.C., Shi, P.Y., Hall, R.A. et al./person-group>. (2008) A highly structured, nuclease-resistant, noncoding RNA produced by flaviviruses is required for pathogenicity. Cell Host Microbe, 4, 579–591.

MacFadden, A., O’Donoghue, Z., Silva, P., Chapman, E.G., Olsthoorn, R.C., Sterken, M.G., Pijlman, G.P., Bredenbeek, P.J. and Kieft, J.S. (2018) Mechanism and structural diversity of exoribonuclease-resistant RNA structures in flaviviral RNAs. Nat Commun, 9, 119.

Chapman, E.G., Moon, S.L., Wilusz, J. and Kieft, J.S. (2014) RNA structures that resist degradation by Xrn1 produce a pathogenic Dengue virus RNA. Elife, 3, e01892.

Chapman, E.G., Costantino, D.A., Rabe, J.L., Moon, S.L., Wilusz, J., Nix, J.C. and Kieft, J.S. (2014) The structural basis of pathogenic subgenomic flavivirus RNA (sfRNA) production. Science, 344, 307–310.

O’Connor, J.P. and Peebles, C.L. (1991) In vivo pre-tRNA processing in Saccharomyces cerevisiae. Mol Cell Biol, 11, 425–439.

Haurwitz, R.E., Jinek, M., Wiedenheft, B., Zhou, K. and Doudna, J.A. (2010) Sequence-and structure-specific RNA processing by a CRISPR endonuclease. Science, 329, 1355–1358.

Knapp, D., Michaels, Y.S., Jamilly, M., Ferry, Q.R.V., Barbosa, H., Milne, T.A. and Fulga, T.A. (2019) Decoupling tRNA promoter and processing activities enables specific Pol-II Cas9 guide RNA expression. Nat Commun, 10, 1490.

Riesenberg, S., Kanis, P., Macak, D., Wollny, D., Dusterhoft, D., Kowalewski, J., Helmbrecht, N., Maricic, T. and Paabo, S. (2023) Efficient high-precision homology-directed repair-dependent genome editing by HDRobust. Nat Methods, 20, 1388–1399.

Yan, J., Oyler-Castrillo, P., Ravisankar, P., Ward, C.C., Levesque, S., Jing, Y., Simpson, D., Zhao, A., Li, H., Yan, W. et al./person-group>. (2024) Improving prime editing with an endogenous small RNA-binding protein. Nature, 628, 639–647.

Ponnienselvan, K., Liu, P., Nyalile, T., Oikemus, S., Maitland, S.A., Lawson, N.D., Luban, J. and Wolfe, S.A. (2023) Reducing the inherent auto-inhibitory interaction within the pegRNA enhances prime editing efficiency. Nucleic Acids Res, 51, 6966–6980.

Zhang, W., Petri, K., Ma, J., Lee, H., Tsai, C.L., Joung, J.K. and Yeh, J.J. (2024) Enhancing CRISPR prime editing by reducing misfolded pegRNA interactions. Elife, 12.

Fonfara, I., Richter, H., Bratovic, M., Le Rhun, A. and Charpentier, E. (2016) The CRISPR-associated DNA-cleaving enzyme Cpf1 also processes precursor CRISPR RNA. Nature, 532, 517–521.

Lieber, M.R. (2010) The mechanism of double-strand DNA break repair by the nonhomologous DNA end-joining pathway. Annu Rev Biochem, 79, 181–211.

Xue, C. and Greene, E.C. (2021) DNA Repair Pathway Choices in CRISPR-Cas9-Mediated Genome Editing. Trends Genet, 37, 639–656.

