## Supplementary material for "An enhanced Eco1 retron editor enables precision genome engineering in human cells without double-strand breaks": Combined Supplemental Materials

**Supplemental Methods**

**qPCR of retron msDNA with Eco1 RTwt and RTmut**

**Plasmids**

Constructs were cloned into the pGEM-3Z plasmid backbone immediately downstream of a T7 promoter.

**Cells**

Huh-7.5 cells were maintained in Dulbecco's modified Eagle's medium (DMEM; Fisher Scientific, catalog no. 11995065) supplemented with 0.1 mM nonessential amino acids (NEAAs; Fisher Scientific, catalog no. 11140076) and 10% fetal bovine serum (FBS; HyClone Laboratories, lot. #AUJ35777).

**Synthesis of retron RT-msr-msd RNAs for qPCR assay to quantify msDNA copy number.**

Ten micrograms of plasmid DNA was linearized by digestion with Not I–HF. Linearized DNA was purified with the MinElute PCR Purification Kit (Qiagen, catalog no. 28004) according to the manufacturer's instructions and diluted to 0.5 µg/µl in elution buffer (EB), and 2 µg was used as a template for in vitro transcription. Transcription was performed using the Ribomax T7 Kit (Promega, catalog no. P1320). The reaction was incubated at 37°C for 30 min followed by 15-min DNase (from kit) treatment at 37°C. To reduce the carryover of residual undigested plasmid or DNA fragments, the DNase-treated RNA was transferred to a new tube before purifying RNA using the RNeasy Mini Kit (Qiagen, catalog no. 74014) following the manufacturer's instructions. The optional on-column DNase digestion step (Qiagen, catalog no. 79254) was included. Capping and polyadenylation were performed using T7 mScript Standard mRNA Production System (CELLSCRIPT, catalog no. C-MSC100625) following the Cap 1 mRNA protocol described in the user's manual, and the RNA was again purified using the RNeasy mini kit without on-column DNase digestion. The expected RNA yield is ~45 to 60 µg per reaction.

**RNA transfection**

Huh-7.5 cells were seeded at $2.5 \times 10^5$ cells per well in six-well plates. The medium was changed to 2 ml of DMEM containing 1.5% FBS and 0.1 mM NEAA just before transfection. For each transfected well (six-well plate), 0.5 µg of retron RNA was mixed with 5 µl of Lipofectamine 2000 (Fisher Scientific, catalog no. 11668019) in 500 µl of Opti-MEM Reduced-Serum Medium (Fisher Scientific, catalog no. 51985034) and incubated at room temperature for 20 min. The mixture was then added to cells and spinoculated by centrifugation at 1000*g* for 30 min at 37°C. Six hours later, the medium was removed and replaced with DMEM containing 10% FBS and 0.1 mM NEAA.

#### Retron msDNA quantification by qPCR

To quantify retron msDNA, 24 h post-transfection, total DNA was extracted from individual wells of 6-well plates using the Qiagen DNeasy Blood & Tissue Kit (Qiagen, catalog no. 69504). 200 ng of total DNA was used as a template together with 1 µl of each primer (3 µM stock) and 10 µl of PowerUp™ SYBR™ Green Master Mix for qPCR (ThermoFisher Scientific; catalog no. A25742). qPCR was performed on QuantStudio 3 with the following forward and reverse primers, respectively: RU-O-32184 5'- TCTGAGTTACTGTCTGTTTTCCTGAAGTC -3'; RU-O-32185
5'- GTCAGAAGAAACGGGTTTCCTCGGCAAG -3'

### sgRNA detection by northern blot with and without Cas9

#### Plasmid transfection and RNA extraction

24 hours before transfection, 1e6 LentiX 293T cells were seeded to a 6-well plate. Cells were transfected the next day using Lipofectamine2000 (Invitrogen) at a 2.5:1 (ul lipo:ug DNA) ratio according to the manufacturer's protocol. Each well was transfected with 1µg of sgBFP plasmid and 1µg of either lentiCas9-Blast or 1µg of GFP. lentiCas9-Blast was a gift from Feng Zhang (Addgene #52962). RNA was extracted by TRIzol according to the manufacturer's protocol 24 hours after plasmid transfection.

#### Northern blotting protocol

First, a 10% acrylamide urea gel was cast (36.4mL DEPC-treated water, 34g Urea, 1.5mL 50X MOPS, 12.6mL 40% acrylamide:bis-acrylamide (37.5:1), 250µL 10% APS, 65µL TEMED) and allowed to solidify at room temperature. Probes were radiolabeled in a 20µL T4 Kinase reaction using 3µL of $\gamma$-32P and purified using a G25 sephadex column. After radiolabeling probes and casting, the acrylamide gel was placed in a BioRad Protean II XI System and filled with 1x MOPS running buffer (80mL of 50X MOPS up to 4L of dH20). 10µg of whole-cell RNA was loaded into each well and run at 300V for about 2-4 hours, until the RiboRuler Low Range Ladder reached the end of the gel. RNA in the gel was transferred to a positively charged nylon membrane using a ThermoFisher OWL HEP Semi-Dry Transfer System, transferring at 300mA for 1 hour at 4ºC. RNA was UV cross linked to the membrane and subsequently stained with 0.02% methylene blue. Membranes were blocked at 65ºC for 1 hour in 9mL of 20X SSC and 21mL of 10% SDS. Radiolabeled probes were then added to blocking solution and temperature was lowered to 42ºC and allowed to hybridize for at least 12 hours.

Each membrane was exposed for 48 hours before imaging (24 hours for loading control probe). Membranes were stripped by 3X 20 minute washes using washing buffer solution (56.25mL 20xSSC, 3.75mL 10%SDS, 315mL DEPC water) at 65ºC. After wash steps, membranes were blocked and hybridized with the probe once again.

**Supplemental Figures**

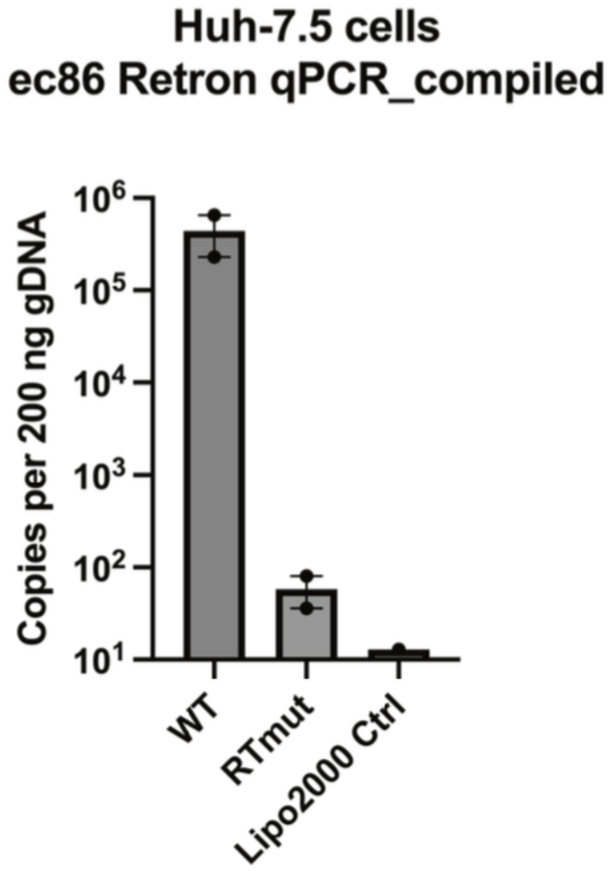

**Supplemental Figure 1. Validation of catalytically inactive Eco1 RTmut by qPCR quantification of retron RT-DNA.** Copies of retron RT-DNA per 200 ng of gDNA for the indicated transfection conditions.

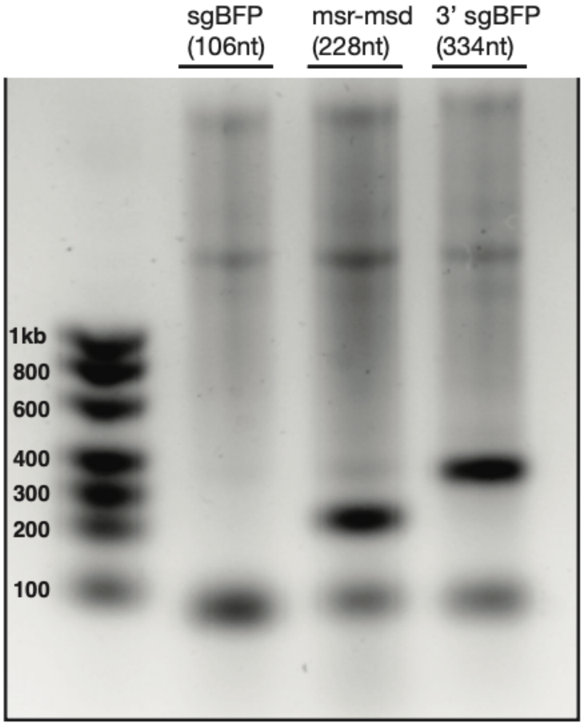

**Supplemental Figure 2. Confirmation of IVT RNA products by agarose gel electrophoresis.** Electrophoretic size separation in agarose gel of sgBFP, msr-msd with internal GFP template, and msr-msd-sgBFP ncRNAs. Each lane was loaded with 200ng of IVT RNA product.

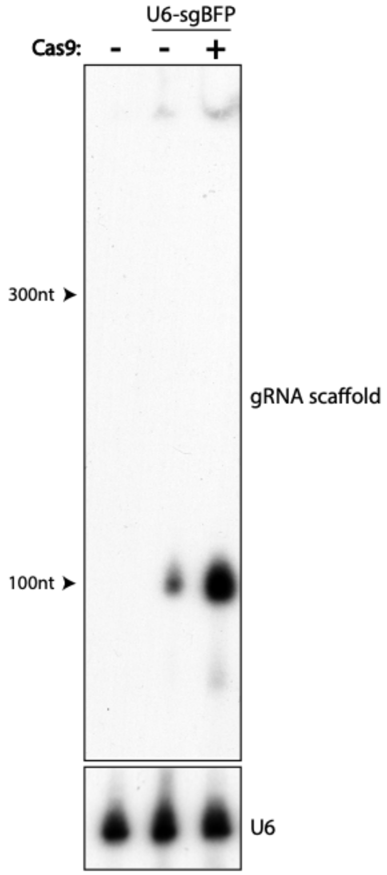

**Supplemental Figure 3. sgRNA stability in the presence and absence of Cas9.** Northern blot analysis of sgRNA from total RNA of HEK293T cells transfected with plasmids expressing sgBFP and either Cas9 or pUC19. $^{32}P$ radiolabeled ssDNA probes were designed to target the scaffold sequence of the sgRNA. Probe sequence is listed in Supplemental Table 1.

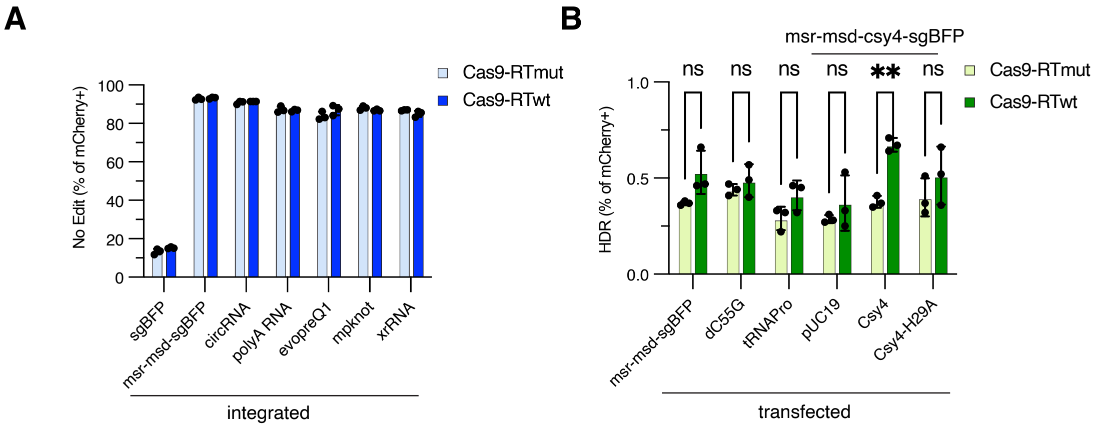

**Supplemental Figure 4. Effect of ncRNA modifications retron editor activity. A)** Analysis of NHEJ or **B)** HDR for the indicated constructs.

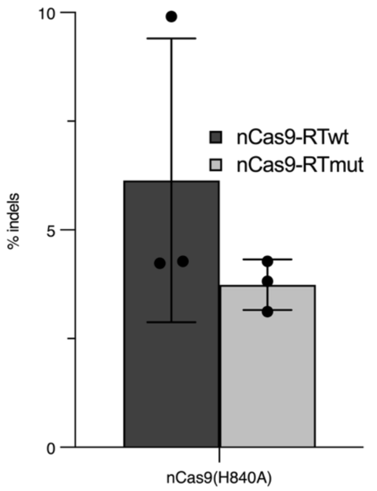

**Supplemental Figure 5. Quantification of NHEJ outcomes at the BFP locus with nCas9(H840A).** Indel frequency at the BFP as a percentage of total reads for HEK293T-BFP cells transfected with optimized xrRNA-evopreQ1-csy4 ncRNA and nCas9(H840A)-RT.

| oligo | type | sequence1 | sequence2 |
|---|---|---|---|
| U6-msr-msd | gBlock | gatgaatactgccatttgtctcttaatt<br>aaagagggcctatttcccatgattcct<br>tcatatttgcatatacgatacaaggct<br>gttagagagataattagaattaatttg<br>actgtaaacacaaagatattagtac<br>aaaatacgtgacgtagaaagtaata<br>atttcttgggtagtttgcagttttaaaatt<br>atgttttaaaatggactatcatatgctt<br>accgtaacttgaaagtatttcgatttct<br>tggctttatatatcttgtggaaaggac<br>gaaacaccgatgcgcacccttagc<br>gagaggtttatcattaaggtcaacct<br>ctggatgttgtttcggcatcctgcattg<br>aatctgagttactgtctgatttcctgaa<br>gtcgtgctgcttcatgtggtcggggta<br>gcggctgaagcactgcacgccgta<br>cgtcagggtggtcacgagggtggg<br>ccagggcaccggcagcttgccgag<br>gaaacccgtttcttctgacgtaaggg<br>tgcgcatttttttaatgatacggcgacc<br>accgagatctacacgaattctttaag<br>cttggcgtaactagatcttttaattaac<br>tgggtcccctcggggttggga | |
| U6-sgBFP | gBlock | gatgaatactgccatttgtctcttaatt<br>aaagagggcctatttcccatgattcct<br>tcatatttgcatatacgatacaaggct<br>gttagagagataattagaattaatttg<br>actgtaaacacaaagatattagtac<br>aaaatacgtgacgtagaaagtaata<br>atttcttgggtagtttgcagttttaaaatt<br>atgttttaaaatggactatcatatgctt<br>accgtaacttgaaagtatttcgatttct<br>tggctttatatatcttgtggaaaggac<br>gaagctgaagcactgcacgccatgt<br>ttaagagctatgctggaaacagcat<br>agcaagtttaaataaggctagtccgt<br>tatcaacttgaaaaagtggcaccga<br>gtcggtgctttttttaatgatacggcga<br>ccaccgagatctacacgaattcttta<br>agcttggcgtaactagatcttttaatta<br>actgggtcccctcggggttggga | |
| U6-msr-msd-sgBFP | gBlock | gatgaatactgccatttgtctcttaatt<br>aaagagggcctatttcccatgattcct<br>tcatatttgcatatacgatacaaggct<br>gttagagagataattagaattaatttg<br>actgtaaacacaaagatattagtac<br>aaaatacgtgacgtagaaagtaata<br>atttcttgggtagtttgcagttttaaaatt<br>atgttttaaaatggactatcatatgctt<br>accgtaacttgaaagtatttcgatttct<br>tggctttatatatcttgtggaaaggac<br>gaaacaccgatgcgcacccttagc<br>gagaggtttatcattaaggtcaacct | aggaaacccgtttcttctgacgtaag<br>ggtgcgcagctgaagcactgcacg<br>ccatgtttaagagctatgctggaaac<br>agcatagcaagtttaaataaggcta<br>gtccgttatcaacttgaaaaagtggc<br>accgagtcggtgctttttttaatgatac<br>ggcgaccaccgagatctacacgaa<br>ttctttaagcttggcgtaactagatcttt<br>taattaactgggtcccctcggggttg<br>gga |

| | | | |
|---|---|---|---|
| | | ctggatgttgtttcggcatcctgcattgaatctgagttactgtctgatttcctgaagtcgtgctgcttcatgtggtcggggtagcggctgaagcactgcacgccgtacgtcagggtggtcacgagggtgggccagggcaccggcagcttgccgaggaaacccgtttcttctgac | |
| U6-sgBFP-msr-msd | gBlock | gatgaatactgccatttgtctcttaattaaagagggcctatttcccatgattccttcatatttgcatatacgatacaaggctgttagagagataattagaattaatttgactgtaaacacaaagatattagtacaaaatacgtgacgtagaaagtaataatttcttgggtagtttgcagttttaaaattatgttttaaaatggactatcatatgcttaccgtaacttgaaagtatttcgatttcttggctttatatatcttgtggaaaggacgaaacaccgctgaagcactgcacgccatgtttaagagctatgctggaaacagcatagcaagtttaaataaggctagtccgttatcaacttgaaaaagtggcaccgagtcggtgcatgcgcacccttagcgagaggtttatcattaaggtcaacctctggatgttgtttcg | tcaacctctggatgttgtttcggcatcctgcattgaatctgagttactgtctgatttcctgaagtcgtgctgcttcatgtggtcggggtagcggctgaagcactgcacgccgtacgtcagggtggtcacgagggtgggccagggcaccggcagcttgccgaggaaacccgtttcttctgacgtaagggtgcgcattttttaatgatacggcgaccaccgagatctacacgaattctttaagcttggcgtaactagatcttttaattaactgggtcccctcggggttggga |
| U6-msr-msd-sgBFPV2 (MCrP3_001) | gBlock | gatgaatactgccatttgtctcttaattaaagagggcctatttcccatgattccttcatatttgcatatacgatacaaggctgttagagagataattagaattaatttgactgtaaacacaaagatattagtacaaaatacgtgacgtagaaagtaataatttcttgggtagtttgcagttttaaaattatgttttaaaatggactatcatatgcttaccgtaacttgaaagtatttcgatttcttggctttatatatcttgtggaaaggacgaaacaccgtgataagattccgtatgcgcacccttagcgagaggtttatcattaaggtcaacctctggatgttgtttcggcatcctgcattgaatctgagttactgtctgatttccttgttggcggcaagctgccggtgccctggcccaccctcgtgaccaccctgacgtacggcgtgcagtgcttcagccgctaccccgaccacatgaagcagcacgacttcccaaccaggaaacccgtttcttctgacg | accaggaaacccgtttcttctgacgtaagggtgcgcatacggaatcttatcacaacaacaacaagctgaagcactgcacgccatgtttaagagctatgctggaaacagcatagcaagtttaaataaggctagtccgttatcaacttgaaaaagtggcaccgagtcggtgcttttttaatgatacggcgaccaccgagatctacacgaattctttaagcttggcgtaactagatcttttaattaactgggtcccctcggggttggga |
| U6-Twister-12bpLinker-msr-msd-sgBFP-12bpLinker-Twister (MCrP3_012) | gBlock | gatgaatactgccatttgtctcttaattaaagagggcctatttcccatgattccttcatatttgcatatacgatacaaggctgttagagagataattagaattaatttgactgtaaacacaaagatattagtacaaaatacgtgacgtagaaagtaataatttcttgggtagtttgcagttttaaaattatgttttaaaatggactatcatatgcttaccgtaacttgaaagtatttcgatttcttggctttatatatcttgtggaaaggacgaaacaccggccatcagtcgccggtcccaagcccggataaaatgggag | aggaaacccgtttcttctgacgtaagggtgcgcagctgaagcactgcacgccatgtttaagagctatgctggaaacagcatagcaagtttaaataaggctagtccgttatcaacttgaaaaagtggcaccgagtcggtgcaaccatgccgactgatggcagaacactgccaatgccggtcccaagcccggataaaagtggagggtacagtccacgcttttttaatgatacggcgaccaccgagatctacacgaattctttaagcttggcgtaactaga |

| | | | |
|---|---|---|---|
| | | ggggcgggaaaccgcctctgccatcagtcggcgtggactgtagatgcgcacccttagcgagaggtttatcattaaggtcaacctctggatgttgtttcggcatcctgcattgaatctgagttactgtctgatttcctgaagtcgtgctgcttcatgtggtcggggtagcggctgaagcactgcacgccgtacgtcagggtggtcacgagggtggccagggcaccggcagcttgccgaggaaacccgtttcttctgacg | tcttttaattaactgggtcccctcggggttggga |
| U6-msr-msd-sgBFP-pA (MCrP3_014) | gBlock | gatgaatactgccatttgtctcttaattaaagagggcctatttcccatgattccttcatatttgcatatacgatacaaggctgttagagagataattagaattaatttgactgtaaacacaaagatattagtacaaaatacgtgacgtagaaagtaataatttcttgggtagtttgcagttttaaaattatgttttaaaatggactatcatatgcttaccgtaacttgaaagtatttcgatttcttggctttatatatcttgtggaaaggacgaaacaccgatgcgcacccttagcgagaggtttatcattaaggtcaacctctggatgttgtttcggcatcctgcattgaatctgagttactgtctgatttcctgaagtcgtgctgcttcatgtggtcggggtagcggctgaagcactgcacgccgtacgtcagggtggtcacgagggtgggccagggcaccggcagcttgccgaggaaacccgtttcttctgacg | aggaaacccgtttcttctgacgtagggtgcgcagctgaagcactgcacgccatgtttaagagctatgctggaaacagcatagcaagtttaaataaggctagtccgttatcaacttgaaaaagtggcaccgagtcggtgcaataaatttttaatgatacggcgaccaccgagatctacacgaattctttaagcttggcgtaactagatcttttaattaactgggtcccctcggggttggga |
| U6-evopreQ1-12bpLinker-msr-msd-sgBFP (MCrP3_006) | gBlock | gatgaatactgccatttgtctcttaattaaagagggcctatttcccatgattccttcatatttgcatatacgatacaaggctgttagagagataattagaattaatttgactgtaaacacaaagatattagtacaaaatacgtgacgtagaaagtaataatttcttgggtagtttgcagttttaaaattatgttttaaaatggactatcatatgcttaccgtaacttgaaagtatttcgatttcttggctttatatatcttgtggaaaggacgaaacaccgTTGACGCGGTTCTATCTAGTTACGCGTTAAACCAACTAGAAAcaacaacaacaaatgcgcacccttagcgagaggtttatcattaaggtcaacctctggatgttgtttcggcatcctgcattgaatctgagttactgtctgatttcctgaagtcgtgctgcttcatgtggtcggggtagcggctgaagcactgcacgccgtacgtcagggtggtcacgagggtgggccagggcaccggcagcttgccgaggaaacccgtttcttctgacg | aggaaacccgtttcttctgacgtagggtgcgcagctgaagcactgcacgccatgtttaagagctatgctggaaacagcatagcaagtttaaataaggctagtccgttatcaacttgaaaaagtggcaccgagtcggtgctttttttaatgatacggcgaccaccgagatctacacgaattctttaagcttggcgtaactagatcttttaattaactgggtcccctcggggttggga |
| U6-mpknot-12bpLinker-msr-msd-sgBFP (MCrP3_008) | gBlock | gatgaatactgccatttgtctcttaattaaagagggcctatttcccatgattccttcatatttgcatatacgatacaaggctgttagagagataattagaattaatttg | aggaaacccgtttcttctgacgtagggtgcgcagctgaagcactgcacgccatgtttaagagctatgctggaaacagcatagcaagtttaaataaggcta |

| | | | |
|---|---|---|---|
| | | actgtaaacacaaagatattagtacaaaatacgtgacgtagaaagtaataatttcttgggtagtttgcagttttaaaattatgttttaaaatggactatcatatgcttaccgtaacttgaaagtatttcgatttcttggctttatatatcttgtggaaaggacgaaacaccgGGGTCAGGAGCCCCCCCCCTGAACCCAGGATAACCCTCAAAGTCGGGGGGCAACCCcaacaacaacaaatgcgcacccttagcgagaggtttatcattaaggtcaacctctggatgttgtttcggcatcctgcattgaatctgagttactgtctgatttcctgaagtcgtgctgcttcatgtggtcggggtagcggctgaagcactgcacgccgtacgtcagggtggtcacgagggtgggccagggcaccggcagcttgccgaggaaacccgtttcttctgacg | gtccgttatcaacttgaaaaagtggcaccgagtcggtgcttttttaatgatacggcgaccaccgagatctacacgaattctttaagcttggcgtaactagatcttttaattaactgggtcccctcggggttggga |
| U6-xrRNA-12bpLinker-msr-msd-sgBFP (MCrP3_010) | gBlock | gatgaatactgccatttgtctcttaattaaagagggcctatttcccatgattccttcatatttgcatatacgatacaaggctgttagagagataattagaattaatttgactgtaaacacaaagatattagtacaaaatacgtgacgtagaaagtaataatttcttgggtagtttgcagttttaaaattatgttttaaaatggactatcatatgcttaccgtaacttgaaagtatttcgatttcttggctttatatatcttgtggaaaggacgaaacaccgTGTCAGGCCTGCTAGTCAGCCACAGTTTGGGGAAAGCTGTGCAGCCTGTAACCCCCCCAGGAGAAGCTGGGAAACCAAGCTcaacaacaacaaatgcgcacccttagcgagaggtttatcattaaggtcaacctctggatgttgtttcggcatcctgcattgaatctgagttactgtctgatttcctgaagtcgtgctgcttcatgtggtcggggtagcggctgaagcactgcacgccgtacgtcagggtggtcacgagggtgggccagggcaccggcagcttgccgaggaaacccgtttcttctgacg | aggaaacccgtttcttctgacgtaagggtgcgcagctgaagcactgcacgccatgtttaagagctatgctggaaacagcatagcaagtttaaataaggctagtccgttatcaacttgaaaaagtggcaccgagtcggtgcttttttaatgatacggcgaccaccgagatctacacgaattctttaagcttggcgtaactagatcttttaattaactgggtcccctcggggttggga |
| U6-xrRNA(C20G)-12bpLinker-msr-msd-sgBFP (MCrP3_010-C20G) | gBlock | gatgaatactgccatttgtctcttaattaaagagggcctatttcccatgattccttcatatttgcatatacgatacaaggctgttagagagataattagaattaatttgactgtaaacacaaagatattagtacaaaatacgtgacgtagaaagtaataatttcttgggtagtttgcagttttaaaattatgttttaaaatggactatcatatgcttaccgtaacttgaaagtatttcgatttcttggctttatatatcttgtggaaaggacgaaacaccgTGTCAGGCCTGCTAGTCAGGCACAGTTTGGGGAAAGCTGTGCAG | aggaaacccgtttcttctgacgtaagggtgcgcagctgaagcactgcacgccatgtttaagagctatgctggaaacagcatagcaagtttaaataaggctagtccgttatcaacttgaaaaagtggcaccgagtcggtgcttttttaatgatacggcgaccaccgagatctacacgaattctttaagcttggcgtaactagatcttttaattaactgggtcccctcggggttggga |

| | | | |
|---|---|---|---|
| | | CCTGTAACCCCCCCAGGAGAAGCTGGGAAACCAAGCTcaacaacaacaaatgcgcacccttagcgagaggtttatcattaaggtcaacctctggatgttgtttcggcatcctgcattgaatctgagttactgtctgatttcctgaagtcgtgctgcttcatgtggtcggggtagcggctgaagcactgcacgccgtacgtcagggtggtcacgagggtgggccagggcaccggcagcttgccgaggaaacccgtttcttctgacg | |
| U6-msr-msd-tRNAPro-sgBFP | gBlock | gatgaatactgccatttgtctcttaattaaagagggcctatttcccatgattccttcatatttgcatatacgatacaaggctgttagagagataattagaattaatttgactgtaaacacaaagatattagtacaaaatacgtgacgtagaaagtaataatttcttgggtagtttgcagttttaaaattatgttttaaaatggactatcatatgcttaccgtaacttgaaagtatttcgatttcttggctttatatatcttgtggaaaggacgaaacaccgtgataagattccgtatgcgcacccttagcgagaggtttatcattaaggtcaacctctggatgttgtttcggcatcctgcattgaatctgagttactgtctgatttccttgttggcggcaagctgccggtgccctggcccaccctcgtgaccaccctgacgtacggcgtgcagtgcttcagccgctaccccgaccacatgaagcagcacgacttcccaaccaggaaacccgtttcttctgacg | accaggaaacccgtttcttctgacgtaagggtgcgcatacggaatcttatcaggctcgttggtctaggggtatgattctcgcttagggtgcgagaggtcccgggttcaaatcccgacgagcccggctgaagcactgcacgccatgtttaagagctatgctggaaacagcatagcaagtttaaataaggctagtccgttatcaacttgaaaaagtggcaccgagtcggtgctttttaatgatacggcgaccaccgagatctacacgaattctttaagcttggcgtaactagatcttttaattaactgggtcccctcggggttggga |
| U6-msr-msd-dC55G-sgBFP | gBlock | gatgaatactgccatttgtctcttaattaaagagggcctatttcccatgattccttcatatttgcatatacgatacaaggctgttagagagataattagaattaatttgactgtaaacacaaagatattagtacaaaatacgtgacgtagaaagtaataatttcttgggtagtttgcagttttaaaattatgttttaaaatggactatcatatgcttaccgtaacttgaaagtatttcgatttcttggctttatatatcttgtggaaaggacgaaacaccgtgataagattccgtatgcgcacccttagcgagaggtttatcattaaggtcaacctctggatgttgtttcggcatcctgcattgaatctgagttactgtctgatttccttgttggcggcaagctgccggtgccctggcccaccctcgtgaccaccctgacgtacggcgtgcagtgcttcagccgctaccccgaccacatgaagcagcacgacttcccaaccaggaaacccgtttcttctgacg | accaggaaacccgtttcttctgacgtaagggtgcgcatacggaatcttatcaggctcgttgggaggtcccgggttgaaatcccggacgagcccgctgaagcactgcacgccatgtttaagagctatgctggaaacagcatagcaagtttaaataaggctagtccgttatcaacttgaaaaagtggcaccgagtcggtgcttttttaatgatacggcgaccaccgagatctacacgaattctttaagcttggcgtaactagatcttttaattaactgggtcccctcggggttggga |
| U6-msr-msd-Csy4-sgBFP | gBlock | | |

| | | |
|---|---|---|
| sgRNA scaffold probe | primer | GTTGATAACGGACTAGCCTTA |
| U6 snoRNA probe | primer | GCCATGCTAATCTTCTCTGTATC |
| T7-msr-msd | gBlock | gatgaatactgccatttgtctcttaattaataatacgactcactatagggatgcgcacccttagcgagaggtttatcattaaggtcaacctctggatgttgtttcggcatcctgcattgaatctgagttactgtctgttttcctgaagtcgtgctgcttcatgtggtcggggtagcggctgaagcactgcacgccgtacgtcagggtggtcacgagggtgggccagggcaccggcagcttgccgaggaaacccgtttcttctgacgtaagggtgcgcagatggccggcatggtcccagcctcctcgctggcgccggctgggcaacaccttcgggtggcgaatgggacttaatgatacggcgaccaccgagatctacacgaattcttaattaactgggtcccctcggggttggga |
| T7-sgBFP | gBlock | gatgaatactgccatttgtctcttaattaataatacgactcactatagctgaagcactgcacgccatgtttaagagctatgctggaaacagcatagcaagtttaaataaggctagtccgttatcaacttgaaaaagtggcaccgagtcggtgcttttgatggccggcatggtcccagcctcctcgctggcgccggctgggcaacaccttcgggtggcgaatgggacttaatgatacggcgaccaccgagatctacacgaattcttaattaactgggtcccctcggggttggga |
| T7-msr-msd-sgBFP | gBlock | gatgaatactgccatttgtctcttaattaataatacgactcactatagggatgcgcacccttagcgagaggtttatcattaaggtcaacctctggatgttgtttcggcatcctgcattgaatctgagttactgtctgttttcctgaagtcgtgctgcttcatgtggtcggggtagcggctgaagcactgcacgccgtacgtcagggtggtcacgagggtgggccagggcaccggcagcttgccgaggaaacccgtttcttctgacgtaagggtgcgcagctgaagcactgcacgccatgtttaagagctatgctggaaacagcatagcaagtttaaataaggctagtccgttatcaacttgaaaaagtggcaccgagtcggtgctttttgatggccggcatggtcccagcctcctcgctggcgccggctgggcaacaccttcgggtggcgaatgggacttaatgatacggcgaccaccgagatctacacgaattcttaattaactgggtcccctcggggttggga |

| | | |
|---|---|---|
| Eco1-RTmut-primer | primer | cctgatctacaccagatatgccaacaatctgacactgagcgcc |
| Eco1-msd-qPCR_F | primer | tctgagttactgtctgatttcctgaagtc |
| Eco1-msd-qPCR_R | primer | gtcagaagaaacgggtttcctcggcaag |

**Supplemental Table 1.** Sequences for all oligos used in this study.
